# miR156-mediated changes in leaf composition lead to altered photosynthetic traits during vegetative phase change

**DOI:** 10.1101/2020.06.22.165977

**Authors:** Erica H. Lawrence, Clint J. Springer, Brent R. Helliker, R. Scott Poethig

## Abstract

- Plant morphology and physiology change with growth and development. Some of these changes are due to change in plant size and some are the result of genetically programmed developmental transitions. In this study we investigate the role of the developmental transition, vegetative phase change (VPC), on morphological and photosynthetic changes.
- We used overexpression of miR156, the master regulator of VPC, to modulate the timing of VPC in *Populus tremula x alba, Zea mays* and *Arabidopsis thaliana* to determine its role in trait variation independent of changes in size and overall age.
- Here we find that juvenile and adult leaves in all three species photosynthesize at different rates and that these differences are due to phase-dependent changes in specific leaf area (SLA) and leaf N but not photosynthetic biochemistry. Further, we found juvenile leaves with high SLA were associated with better photosynthetic performance at low light levels.
- This study establishes a role for VPC in leaf composition and photosynthetic performance across diverse species and environments. Variation in leaf traits due to VPC are likely to provide distinct benefits under specific environments and, as a result, selection on the timing of this transition could be a mechanism for environmental adaptation.

## Introduction

As plants age they go through developmental transitions that impact their form and function. One of these transitions occurs as plants shift between juvenile and adult vegetative growth phases. This developmental transition is known as vegetative phase change (VPC) and has been observed across phylogenetically diverse groups of plants from mosses to angiosperms. This transition is controlled by expression levels of the highly conserved microRNA, miR156 and in some species the closely related miR157 (Willmann & Poethig, 2007; Axtell & Bowman, 2008; Zhang *et al*., 2015). miR156/7 are expressed at high levels in leaves produced early in development, and negatively regulate the expression of their targets, the Squamosa Promoter Binding Protein-Like (SPL) transcription factors. Expression of miR156/7 declines later in development, alleviating the transcriptional and translational repression of these SPL genes. This increase in SPL expression promotes adult vegetative traits, leading to vegetative phase change (Wu & Poethig, 2006; Wu *et al*., 2009; Wang *et al*., 2011; Xu *et al*., 2016; He *et al*., 2018). The traits that change during VPC are species-dependent, but broadly include changes in leaf morphology, growth rate, growth form, and reproductive competence (Poethig, 1990; Bassiri *et al*., 1992; Bongard-Pierce *et al*., 1996; Telfer *et al*., 1997; Wang *et al*., 2011; Feng *et al*., 2016; Leichty & Poethig, 2019; Silva *et al*., 2019).

How plants respond to dynamic challenges in their environment varies with age (Cavender-Bares & Bazzaz, 2000; Niinemets, 2010; Kitajima *et al*., 2013; Hahn & Orrock, 2016) and has important implications for plant community composition, the competitive ability of different species, and their response to future climate change (Parish & Bazzaz, 1985; Lamb & Cahill, 2006; Moll & Brown, 2008; Piao *et al*., 2013; Spasojevic *et al*., 2014; Kerr *et al*., 2015; Lasky *et al*., 2015). For example, seedlings are particularly vulnerable to factors such as shading, drought, disturbance and herbivory (Kabrick *et al*., 2015; Charles *et al*., 2018) and often experience a high rate of mortality (Grossnickle, 2012). Species that are able to transition to a more resilient phase for their environment are likely to have a competitive advantage. Although it is reasonable to assume that VPC plays an important role in this process, how VPC affects the response of plants to various biotic and abiotic stresses is still poorly understood.

Defining the role of VPC in plant physiology is difficult because this transition occurs concurrently with changes in plant size and age. Juvenile leaves and branches are produced on smaller plants than adult organs, and thus are exposed to different amounts and types of endogenous factors (e.g. hormones, carbohydrates). Furthermore, light, temperature, and humidity vary according to position within the canopy and proximity to the ground (Evans & Coombe, 1959; Waggoner & Reifsnyder, 1968; Shuttleworth *et al*., 1985; Canham, 1988; Canham *et al*., 1994; Still *et al*., 2019), meaning that juvenile and adult organs exist in different microclimates. Finally, the temporal separation between the production of juvenile and adult organs means that seasonal changes in environmental conditions also contribute to age-dependent differences in physiological traits. These problems can only be addressed by varying the timing of VPC under controlled environmental conditions, independent of shoot growth.

The importance of various leaf traits—including photosynthetic traits, specific leaf area, leaf nitrogen content, and gas exchange—for plant growth and survival have been well documented (Lusk & Del Pozo, 2002; Poorter & Bongers, 2006; Modrzynski *et al*., 2015). Previous studies have shown that photosynthetic genes are differentially expressed in juvenile and adult maize leaves (Strable *et al*., 2008; Beydler, 2014), but comparisons of various photosynthetic traits in these leaves have produced inconclusive and sometimes conflicting results (Bond, 2000; Steppe *et al*., 2011; Kuusk *et al*., 2018a, b; Sun *et al*., 2018). The basis for these inconsistencies is unclear, but the compounding effects of variation in plant size, leaf age, environment, and time of year are possibilities (Bauer & Bauer, 1980; Bond, 2000; Ishida *et al*., 2005; Velikova *et al*., 2008; Steppe *et al*., 2011). Although these effects can be minimized through techniques such as grafting, *in vitro* rejuvenation, and pruning (Hutchison *et al*., 1990; Huang *et al*., 2003; Kubien *et al*., 2007; Jaya *et al*., 2010), these methods do not completely distinguish the effect of vegetative phase change from other factors that may contribute to these differences. For example, grafting old shoots to young roots is often used to determine if a trait is dependent on plant size. However, miR156 is a mobile microRNA and can move across a graft junction (Marin-Gonzalez & Suarez-Lopez, 2012; Bhogale *et al*., 2014; Fouracre & Poethig, 2019; Ahsan *et al*., 2019), so this approach does not necessarily eliminate the effect of this key regulator of vegetative identity. Similarly, the methods that are typically used to induce vegetative rejuvenation (*in vitro* culture, pruning) affect both the level of miR156 (Irish & Karlen, 1998; Li *et al*., 2012) and plant size.

We used overexpression of miR156 in three species—*Populus tremula x alba, Zea mays*, and *Arabidopsis thaliana*—to delay the timing of vegetative phase change, allowing us to differentiate traits associated with this developmental transition from those regulated by plant size or age. Our results demonstrate that juvenile leaves are photosynthetically distinct from adult leaves, and that this difference can be attributed primarily to the morphological differences between these leaves, not to a fundamental difference in biochemistry of photosynthesis.

## Materials and Methods

### Plant material

*Populus tremula x alba* line 717□1B4 and two independent miR156 overexpressor lines, 40 and 78, described in Lawrence *et al*., (2020) were obtained by *in vitro* propagation and hardened on propagation media as described in Meilan & Ma (2006). Plants were then transplanted to Fafard 2 growing mix (Sangro Horticulture, Massachusetts, USA) in 0.3 L pots in the greenhouse at the University of Pennsylvania (39.9493°N, 75.1995°W, 22.38 m a.s.l.) and kept in plastic bags for increased humidity for 2 weeks. Plants were transferred to 4.ML pots with Fafard 52 growing mix 3 weeks later and fertilized with Osmocote classic 14Ū14Ū14 (The Scotts Company, Marysville, OH, USA). Plants were additionally fertilized once a week with Peters 20□ 10□20 (ICL Fertilizers, Dublin, OH, USA). Greenhouse conditions consisted of a 16□hr photoperiod with temperatures between 22 and 27°C. Light levels were based on natural light and supplemented with 400 □ W metal halide lamps (P.L. Light Systems, Ontario, Canada) with daily irradiances of 300 to 1,500 μmol m^-2^ s^-1^. All settings controlled by Priva (Ontario, Canada) and Microgrow (Temecula, Canada) greenhouse systems.

*Populus tremula x alba* seeds from Sheffield’s Seed Company (Locke, NY) were germinated on a layer of vermiculite on top of Fafard-2 growing mix in 0.64-L pots in the greenhouse under conditions described above. Seedlings were transplanted to 1.76-L pots with Fafard-52 growing mix with Osmocote classic 14-14-14 one month after germination and were then transplanted to 4.2-L pots 3 months following the previous transplant.

*Zea mays* seeds with the *Corngrass 1 (Cg1)* mutation (stock 310D)—which consists of a tandem duplication of miR156b/c primary sequences described in Chuck et al. (2007)— and W22 inbred lines were obtained from the Maize Genetics Cooperation Stock Center (Urbana, IL). Plants heterozygous for *Cg1* were crossed to W22 to produce the *Cg1/+ and +/+* siblings used in this study. Seeds were planted in 9.09-L pots with Fafard-52 growing mix and fertilized with Osmocote classic 14-14-14 in the greenhouse under growing conditions described above.

*Arabidopsis thaliana* of the Col genetic background and 35S:miR156 overexpressor mutants described in Wu & Poethig (2006) were planted in 0.06-L pots with Fafard-2 growing mix as described by Flexas et al. (2007). Beneficial nematodes (*Steinernema feltiae*, BioLogic, Willow Hill, PA), Marathon^®^ 1 % granular insecticide and diatomaceous earth were added to the growing mix for better plant growth. Planted seeds were placed at 4°C for 3 days before being grown at 22°C in Conviron growth chambers under short days (10 hrs. light/14 hrs. dark) at 60 μmol m^-2^ s^-1^ light to obtain leaves large enough to fit in the gas exchange chamber. Plants were fertilized with Peters 20-10-20 every other week.

Individuals from genotypes of all species were positioned in a randomized fashion in the greenhouse and rotated frequently. Planting was staggered across two, three and five months for *Arabidopsis, P. tremula x alba* and *Z. mays* respectively.

### Leaf samples

All measurements and samples were conducted on the uppermost fully expanded leaf. In *P. tremula x alba* 717-1B4 and miR156 overexpressor lines leaves 10, 15, 20 and 25 were measured. Leaves 10 and 15 in the wild-type 717-1B4 line were juvenile and leaves 20 and 25 were adult as determined by petiole shape and abaxial trichome density as described in Lawrence *et al*., (2020). All measured leaves in the miR156 overexpressor lines were juvenile. In the Poplar plants germinated from seed, leaves 1-52 were measured with a transition to adult between leaf 20 and 30 as determined via petiole shape and trichome density. In *Z. mays*, leaves 2-11 were measured with leaves 1-5 juvenile in wild-type plants and all leaves juvenile in *Cg1* mutants. Developmental stage in maize was determined by the presence or absence of epicuticular wax and trichomes as described in Poethig (1988). In *A. thaliana* leaves 5 and 10 were measured where leaf 5 was juvenile and 10 was adult in wild-type plants, as determined by the presence or absence of abaxial trichomes, and all leaves were juvenile in miR156 overexpressors.

Throughout this manuscript “juvenile” and “adult” leaves refer to those naturally juvenile or adult in the wild-type lines and “juvenilized” leaves refer to those leaves of juvenile phenotype in the miR156 overexpressor lines located at leaf positions that would normally be adult.

### Gas exchange measurements

All gas exchanges measurements were made using a Li-6400 portable photosynthesis machine (Li-Cor Environmental) at a leaf temperature of 25°C following acclimatization to starting chamber conditions. Photosynthetic capacity in *A. thaliana* was measured using steady-state AC_i_ curves measuring A_net_ at reference [CO_2_] of 400, 200, 50, 100, 150, 200, 250, 300, 600, 800, 1000, and 1200 ppm, at a flow rate of 300 μmol air sec^-1^, minimum wait time of 2 mins, and light level of 1000 μmol m^-2^ s^-1^. *Z. mays* AC_i_ curves measured A_net_ at reference [CO_2_] of 400, 350, 300, 250, 200, 150, 100, 50, 400, 500, 600, 700, 800, 1000, 1200 ppm, at a flow rate of 400 μmol air sec^-1^, minimum wait time of 2 mins, and light level of 1800 μmol m^-2^ s^-1^.

Photosynthetic capacity in *P. tremula x alba* was measured using Rapid ACi Response (RACiR) curves as described in Lawrence, Stinziano, and Hanson (2019). Briefly, A_net_ was measured from reference [CO_2_] of 300 to 800 μmol m^-2^ s^-1^ at 60 μmol mol^-1^ min^-1^ CO_2_ and a light level of 1500 μmol m^-2^ s^-1^. This technique was used to expedite measurements after development of the RACiR technique for the Li-6400 showed no significant differences from steady-state ACi curves.

Light response curves were performed in all three species at a reference [CO_2_] of 400 ppm. A_net_ was measured at light levels of 1000, 800, 600, 300, 200, 150, 100, 75, 50, 25, 0 μmol m^-2^ s^-1^ in *A. thaliana*, 1800, 1500, 1200, 1000, 800, 600, 300, 200, 150, 100, 75, 50, 25, 0 μmol m^-2^ s^-1^ in *Z. mays* and 1500, 1200, 1000, 800, 600, 300, 200, 150, 100, 75, 50, 25, 10 and 0 μmol m^-2^ s^-1^ in *P. tremula x alba*. Flow rate, leaf temperature and minimum wait times were the same as for ACi curves.

Low light photosynthetic rates depicted in figure 5 were obtained by averaging photosynthetic rates over a 2 min period at light levels approximately 2-3x the light compensation point. These values were 25 μmol m^-2^ s^-1^ in *P. tremula x alba* and *A. thaliana* and 50 μmol m^-2^ s^-1^ in *Z. mays*. All leaves were acclimated to the chamber conditions before measurements began and flow rate and leaf temperature were consistent with previously described measurements.

Daytime respiration rates were determined by averaging A_net_ at 0 μmol m^-2^ s^-1^ irradiance over a one-minute period after the leaves were dark adapted for 1 hour.

### Leaf Fluorescence

Light and dark-adapted fluorescence was determined using a Li-6400 equipped with fluorometer head. Light adapted measurements were taken using a multiphase flash with a 250 ms phase 1, 500 ms phase 2 with a 20% declining ramp and 250 ms phase 3 after leaves acclimated to saturating light values of 1000, 1800, and 1500 μmol m^-2^ s^-1^ for *A. thaliana, Z. mays* and *P. tremula x alba* respectively. Dark-adapted fluorescence measurements were taken using an 800 ms saturating rectangular flash after dark adapting leaves for 1 hour.

### Leaf nitrogen, chlorophyll and specific leaf area

Leaf tissue was sampled after gas exchange; one subsample for each leaf was dried at 60°C until constant mass to determine SLA. Dried tissues were ground using a mortar and pestle. Leaf nitrogen was measured in the dried samples using an ECS 4010 CHNSO Analyzer (Costech Analytical Technologies INC, Valencia, CA, USA). A second subsample was frozen and used for chlorophyll quantification. Chlorophyll was extracted using 80% acetone and quantified using a spectrophotometer according to equations found in Porra, Thompson, and Kriedemann (1989).

### Leaf cross sections

Fresh leaf tissue from the middle of fully expanded leaves at positions 5 and 10 of *A. thaliana*, 10 and 25 of *P. tremula x alba* and 4 and 11 of *Z. mays* in both wild-type and miR156 overexpressor lines was cut and fixed with a 10x FPGA solution overnight. Samples were then washed with 50% ethanol and dehydrated through an ethanol/*t*-butyl alcohol (TBA) series with 2 hour incubations at room temperature for each step. Sections in 100% TBA were subsequently transferred to Paraplast plus embedding medium at 60°C and incubated for 48 hours. Embedded samples were set in molds and cut into 12μm sections using a microtome. Samples were floated on 0.01% Sta-on on glass slides and dried at 40°C. Samples were then deparaffinized in xylenes and rehydrated through an ethanol series for staining with 1% Safranin O in 50% ethanol and subsequent dehydrating for staining with 0.1% Fast green in 95% ethanol. Once fully stained and dehydrated, sections were mounted in permount and visualized and photographed using an Olympus BX51 light microscope and DP71 digital camera.

### Curve fitting

The {plantecophys} package in Duursma (2015) was used for fitting AC_i_ curves to determine V_cmax_ and J_max_ using the bilinear function for *A. thaliana* and *P. tremula x abla*. The C_4_ photosynthesis estimation tool presented in Zhou, Akçay, and Helliker (2019) based on Yin et al. (2011) was used for fitting AC_i_ curves for *Z. mays*.

Light response curves were analyzed using the {AQ Curve fitting} script in R (Tomeo, 2019) which uses equations based on a standard non-rectangular hyperbola model fit described in Lobo et al. (2013).

### Data analysis

All statistical analyses were performed in JMP ^®^ Pro v. 14.0.0 (SAS Institute Inc., Cary, NC). Gas exchange and leaf composition traits between adult, juvenile and juvenilized leaves were compared by one-way ANOVA and a student’s *t* test (α = 0.05) where developmental stage was the main effect. Traits were considered to be affected by developmental phase when adult leaves were significantly different from both juvenile and juvenilized leaves with the same trend. The effect of leaf position on measured traits was determined by two-way ANOVA with leaf position and genotype as the main effects. Because developmental phase and leaf position are coordinated in wild-type plants, many traits affected by development showed significant leaf position effects (*p* < 0.05). Of these traits, those that showed no significant interaction between leaf position and genotype, where there were no significant differences between wild-type and miR156 overexpressor plants that do not produce adult leaves, are affected by leaf position independent of leaf developmental stage. Photosynthetic nitrogen use efficiency was determined using least squares linear regression analysis across all leaves and was compared by ANCOVA with developmental stage as the covariate.

## Results

### Photosynthetic rates differ between juvenile and adult leaves

The rate of light-saturated area-based photosynthesis (A_max_ Area) was significantly different in juvenile and adult leaves of *P. tremula x alba* and *A. thaliana*, but was not significantly different in maize (Fig. **1**, Table 1). In *P. tremula x alba*, adult leaves had a 26% greater A_max_ Area compared to their juvenile counter parts, whereas in *A. thaliana*, adult leaves had a 57% greater A_max_ Area than juvenile leaves. The phase-dependence of this difference was confirmed by the phenotype of lines over-expressing miR156. In *P. tremula x alba*, the A_max_ Area of adult leaves was, respectively, 104% and 105% greater than the A_max_ Area of the corresponding juvenilized leaves in lines 40 and 78, whereas in Arabidopsis, the A_max_ Area of adult leaves was 42% higher than that of juvenilized leaves.

**Figure 1.**
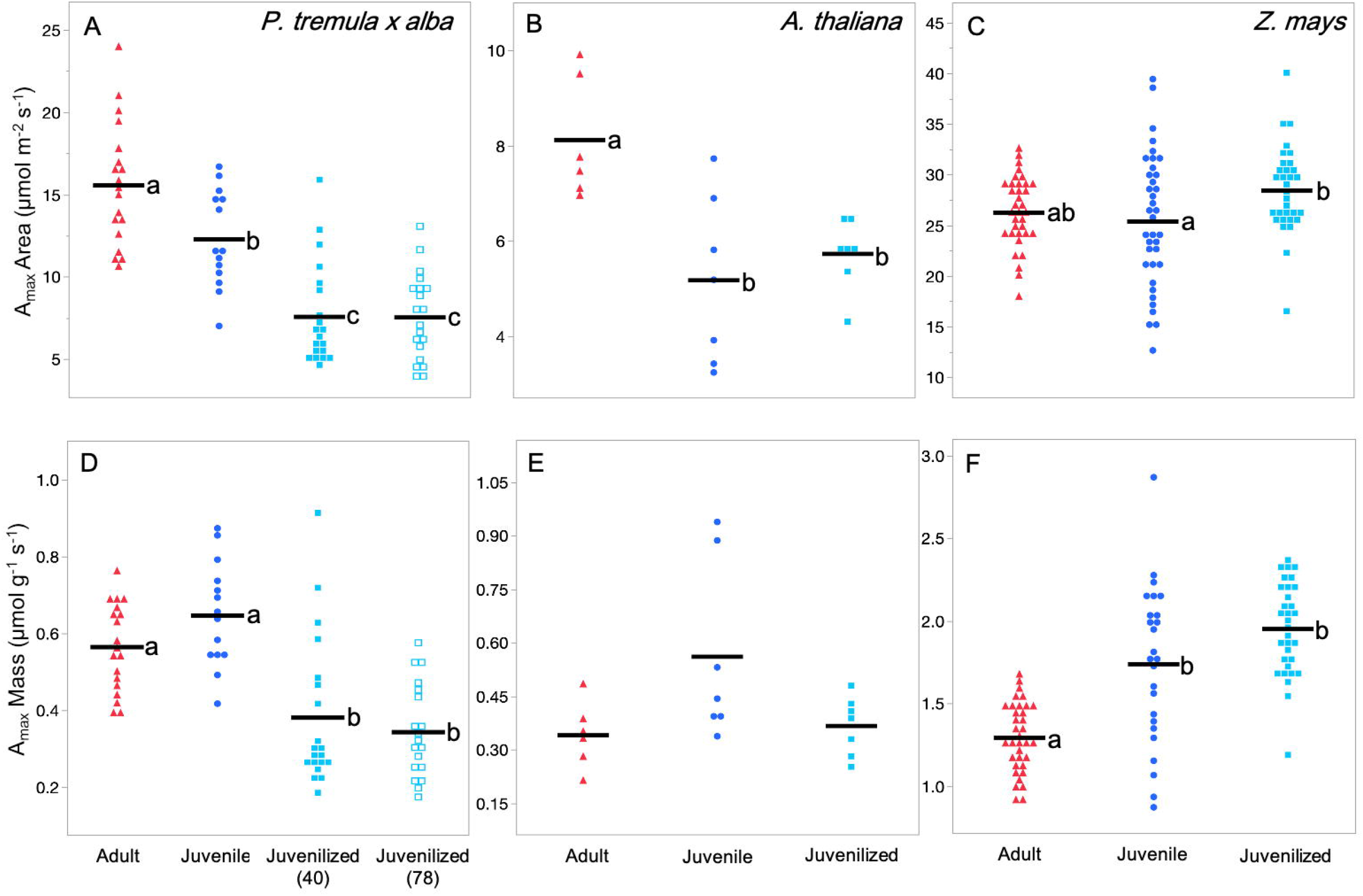
Photosynthetic rates of adult (red ▲), juvenile (blue ●) and juvenilized (light blue ▀ or □) leaves of *P. tremula x alba* (A, D), *A. thaliana* (B, E) and *Z. mays* (C, F). Traits depicted are area-based maximum net photosynthetic rate (A-C, A_max_ Area) and mass-based maximum net photosynthetic rate (D-F, A_max_ mass). Means presented as black horizontal lines. Different lowercase letters indicate means of developmental stage are significantly different according to Student’s *T*(*P* < 0.05).

**Table 1.**
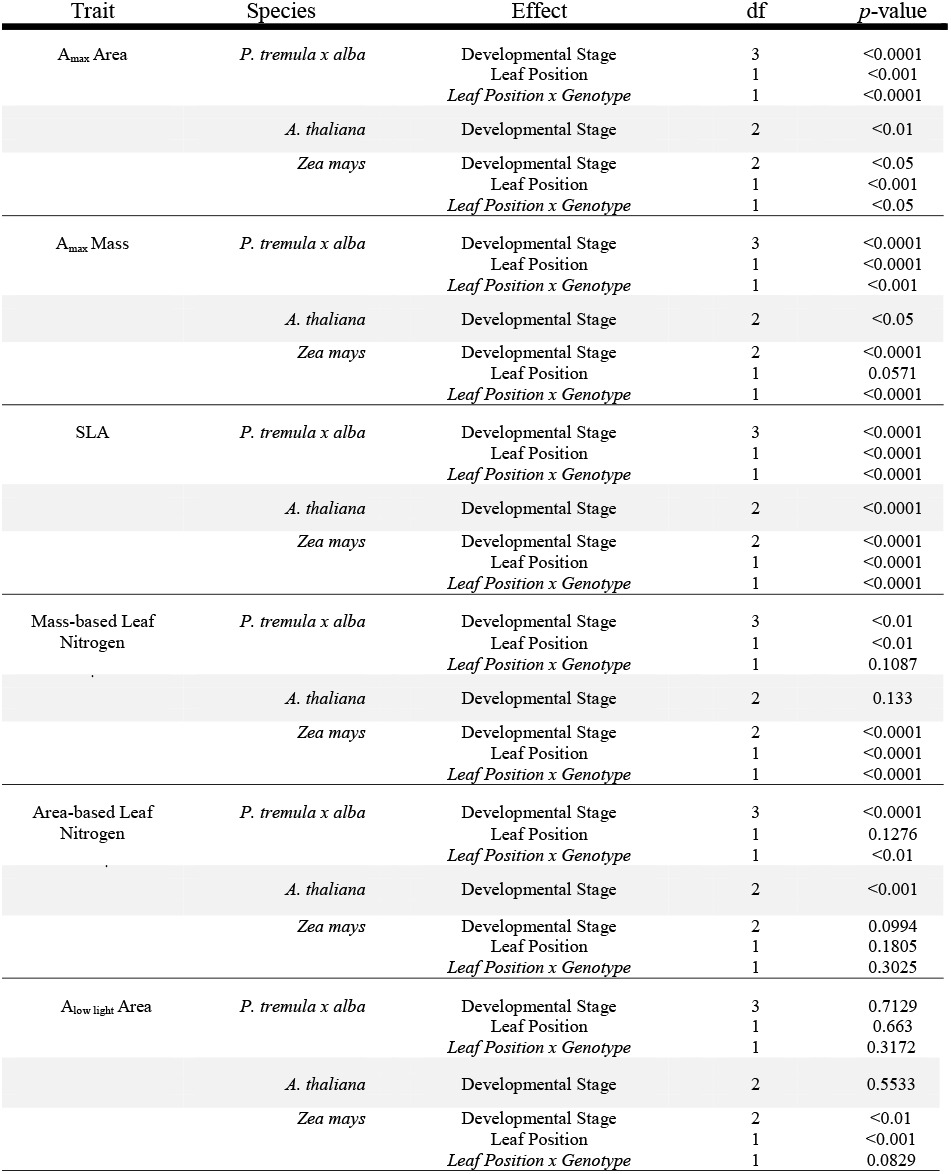
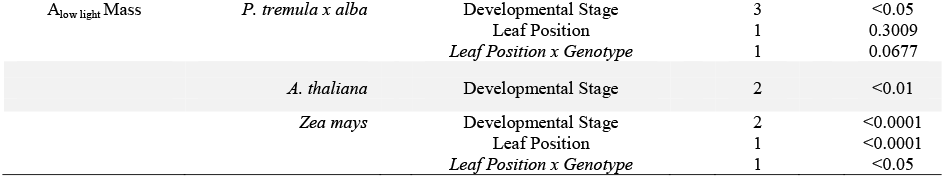
Statistical results for leaf traits depicted in figures 1, 2 and 4. *P*-values determined by one-way ANOVA with developmental stage as the effect variable and two-way ANOVA with leaf position and genotype as the effect variables. Developmental stages are adult, juvenile and juvenilized; genotypes are wild-type and miR156 overexpressors and leaf positions are 2-11 in *Z. mays* and 10, 15, 20 and 25 in *P. tremula x alba*. Leaf position is shown to have an effect on a trait independent of developmental stage when *p* < 0.05 for Leaf position but not for Leaf position x Genotype.

Mass-based photosynthetic rates (A_max_ Mass) were lower in adult leaves than in juvenile leaves in all three species, although this difference was only statistically significant in maize (Fig. **1**, Table 1). In maize juvenilized leaves had essentially the same A_max_ Mass as normal juvenile leaves, suggesting that the difference in A_max_ Mass between juvenile and adult leaves is phase-dependent. However, in *P. tremula x alba* and *A. thaliana*, the A_max_ Mass of juvenilized leaves was significantly lower than that of juvenile leaves, and was more similar to that of adult leaves.

### Leaf morphology and composition is phase-dependent

Inconsistencies in the relationship between leaf identity and A_max_ on an area or mass basis across species suggests that leaf-to-leaf variation in the rate of photosynthesis is either determined by variation in the leaf area/mass relationship or by variation in the photosynthetic biochemistry in these species. *P. tremula x alba* and *A. thaliana* both undergo C_3_ photosynthesis whereas maize is a C4 plant, so it is reasonable to assume that the factors contributing to developmental variation in photosynthesis in these species could be quite different. To address this issue, we measured morphological, chemical, and physiological traits in adult, juvenile, and juvenilized leaves of these species.

Specific leaf area (SLA) represents the amount of area per unit of leaf mass, and is a proxy for the thickness or density of the leaf blade; in general, leaves with a high SLA are thinner than leaves with a low SLA. Adult leaves of all three species had a significantly lower SLA than juvenile leaves (Fig. **2A-C**, Table 1). Furthermore, the SLA of juvenilized leaves was significantly higher than that of adult leaves, and was similar, if not identical to, the SLA of juvenile leaves in both *P. tremula x alba* and maize. This result suggests that SLA is phase dependent in all three species.

**Figure 2.**
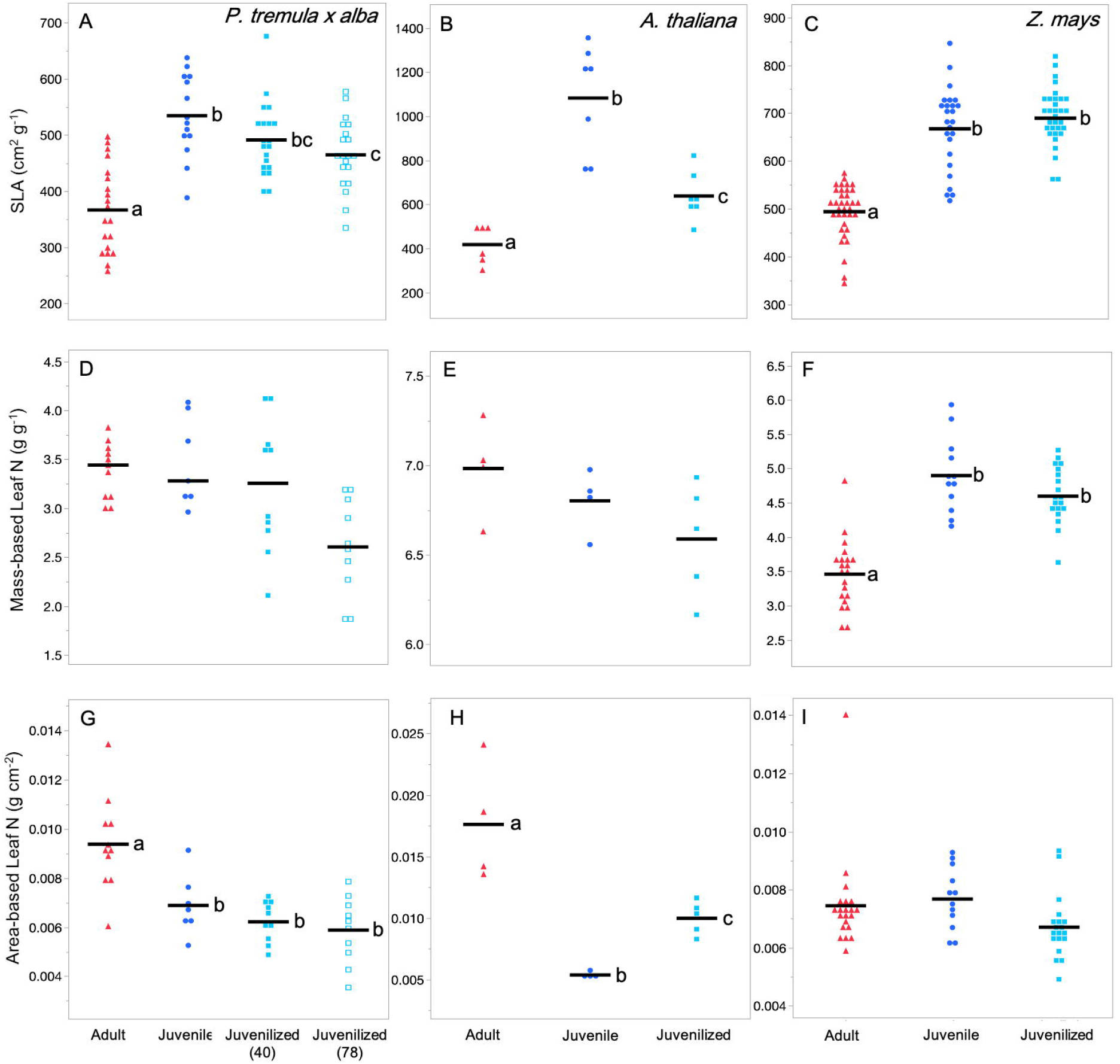
Leaf morphological and compositional traits of adult, juvenile and juvenilized leaves of *P. tremula x alba* (A, D, G), *A. thaliana* (B, E, H) and *Z. mays* (C, F, I). Traits depicted are specific leaf area (SLA, A-C), mass-based leaf Nitrogen content (D-F) and area-based leaf Nitrogen content (G-I). Lettering and symbols are the same as Figure 1.

The relationship between leaf nitrogen (leaf N) and phase identity varied depending on whether this trait was measured on an area or mass basis, and was similar to the results obtained for photosynthetic rates. Measured on a mass basis, leaf N was not significantly different in juvenile and adult leaves of *P. tremula x alba* or *A. thaliana*, and was not significantly different between juvenilized and adult leaves of these species. However, in maize, leaf N/mass was significantly lower in adult leaves than in either juvenile or juvenilized leaves. Thus, leaf N/mass is a phase dependent trait in maize, but not in *P. tremula x alba* or *A. thaliana*. The opposite result was obtained when leaf N was measured as a function of leaf area. In both *P. tremula x alba* and *A. thaliana*, leaf N/area was significantly higher in adult leaves than in juvenile or juvenilized leaves, implying that it phase-dependent in these species. However, there was no significant difference in the leaf N/area of adult, juvenile, or juvenilized leaves in maize (Fig. **2D-I**, Table 1).

SLA and leaf N were significantly correlated with phase-dependent photosynthetic rates (A_max_ Area in *P. tremula x alba* and *A. thaliana;* A_max_ Mass in maize) in all three species (Fig. **3**). SLA was negatively correlated with A_max_ Area in *P. tremula x alba* and *A. thaliana* and positively correlated with A_max_ Mass in *Z. mays*. Leaf N is positively correlated with A_max_ Area in *P. tremula x alba* and *A. thaliana* and A_max_ Mass in *Z. mays*. However, photosynthetic nitrogen use efficiency (PNUE), calculated as the relationship between A_max_ and leaf N, did not vary based on leaf developmental phase (Table 3).

**Figure 3.**
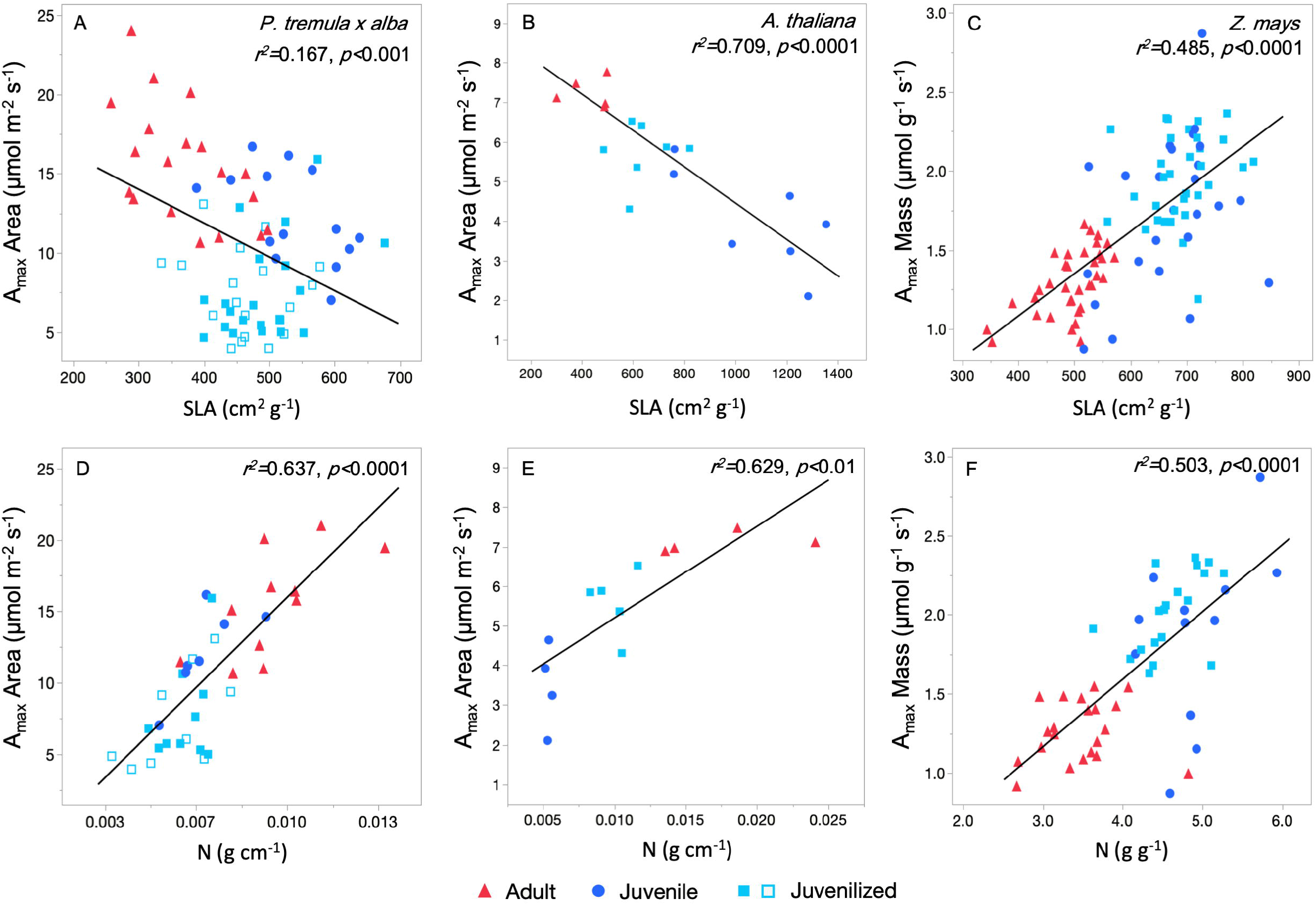
Phase-dependent photosynthetic rates are significantly correlated with leaf composition traits in *P. tremula x alba* (A, D), *A. thaliana* (B, E) and *Z. mays* (C, F). *P. tremula x alba* and *A. thaliana* show significant differences between developmental phase in area-based measures whereas *Z. mays* shows phase-dependence in mass-based measures. Symbols are the same as Figure 1. Linear fit for panel A) A_max_ Area = 20.41 – 0.02134(SLA), B) A_max_ Area = 9.049-0.004591(SLA), C) A_max_ Mass = 0.01188 + 0.002678(SLA), D) A_max_ Area = −2.948 + 1911(N area), E) A_max_ Area = 2.867 + 232.4(N area), F) A_max_ Mass = −0.1081 + 0.4253(N mass).

We also compared Chlorophyll *a* and *b* (Chl_a+b_) levels and ratios between adult, juvenile and juvenilized leaves. Chl_a+b_ was not significantly different across leaves of different developmental phases however, the ratio between Chl_*a*_ and Chl_*b*_ (Chl a:b ratio) was phase-dependent in all three test species (Table 2). Changes in Chl a:b ratios followed the same trends as Leaf N with lower ratios in juvenile and juvenilized leaves than adult leaves of *A. thaliana* and *P. tremula x alba* and the opposite in *Z. mays*. As Chla is associated with more proteins than Chlb, these data support one another.

**Table 2.** Additional leaf traits for adult, juvenile and juvenilized leaves of *P. tremula x alba, A. thaliana* and *Zea mays. P*-values determined by one-way ANOVA with developmental stage as the effect variable. Student’s *T*-test was conducted on traits where *p* < 0.05, means significantly different from each other depicted by different lower-case letters.

| Trait | Species | Developmental Stage | Mean ± SE | N | df | *p*-value |
| --- | --- | --- | --- | --- | --- | --- |
| Chl<sub>a+b</sub><br>($\mu\text{g mg}^{-1}$ FW) | *P. tremula x alba* | Adult | 12.35 ± 4.27 | 20 | 3 | 0.4278 |
| | | Juvenile | 15.02 ± 5.3 | 13 | | |
| | | Juvenilized-40 | 19.27 ± 4.27 | 20 | | |
| | | Juvenilized-78 | 21.72 ± 4.27 | 20 | | |
| | *A. thaliana* | Adult | 1.65 ± 0.99 | 6 | 2 | 0.2728 |
| | | Juvenile | 2.23 ± 0.99 | 6 | | |
| | | Juvenilized | 3.93 ± 0.99 | 5 | | |
| | *Zea mays* | Adult | 33.05 ± 8.98 | 30 | 2 | 0.1439 |
| | | Juvenile | 28.84 ± 8.44 | 34 | | |
| | | Juvenilized | 51.69 ± 8.70 | 32 | | |
| Chl a:b ratio | *P. tremula x alba* | Adult | 1.17 ± 0.04 a | 20 | 3 | <0.0001 |
| | | Juvenile | 0.93 ± 0.05 b | 13 | | |
| | | Juvenilized-40 | 0.99 ± 0.04 b | 20 | | |
| | | Juvenilized-78 | 0.99 ± 0.04 b | 20 | | |
| | *A. thaliana* | Adult | 1.76 ± 0.09 a | 6 | 2 | <0.05 |
| | | Juvenile | 1.47 ± 0.09 b | 6 | | |
| | | Juvenilized | 1.46 ± 0.09 b | 5 | | |
| | *Zea mays* | Adult | 0.97 ± 0.05 a | 30 | 2 | <0.01 |
| | | Juvenile | 1.24 ± 0.05 b | 34 | | |
| | | Juvenilized | 1.20 ± 0.05 b | 32 | | |
| $V_{\text{cmax}}$<br>($\mu\text{mol m}^{-2} \text{s}^{-1}$) | *P. tremula x alba* | Adult | 44.07 ± 4.69 | 15 | 3 | 0.1136 |
| | | Juvenile | 39.17 ± 5.24 | 12 | | |
| | | Juvenilized-40 | 40.16 ± 5.24 | 12 | | |
| | | Juvenilized-78 | 27.84 ± 4.85 | 14 | | |
| | *A. thaliana* | Adult | 42.11 ± 6.21 | 7 | 2 | 0.7371 |
| | | Juvenile | 48.47 ± 6.21 | 7 | | |
| | | Juvenilized | 42.80 ± 6.71 | 6 | | |
| | *Zea mays* | Adult | 35.45 ± 3.42 | 5 | 1 | 0.0586 |
| | | Juvenilized | 25.42 ± 3.12 | 5 | | |
| $J_{\text{max}}$<br>($\mu\text{mol m}^{-2} \text{s}^{-1}$) | *P. tremula x alba* | Adult | 68.81 ± 6.95 a | 15 | 3 | <0.05 |
| | | Juvenile | 66.17 ± 7.76 ab | 12 | | |
| | | Juvenilized-40 | 62.33 ± 7.76 ab | 12 | | |
| | | Juvenilized-78 | 41.65 ± 7.19 b | 14 | | |
| | *A. thaliana* | Adult | 94.55 ± 16.18 | 7 | 2 | 0.676 |
| | | Juvenile | 113.23 ± 16.18 | 7 | | |
| | | Juvenilized | 96.20 ± 17.48 | 6 | | |
| | *Zea mays* | Adult | 204.29 ± 23.24 | 5 | 1 | 0.0507 |
| | | Juvenilized | 133.36 ± 21.22 | 5 | | |
| $\Phi$ | *P. tremula x alba* | Adult | 0.06 ± 0.004 | 19 | 3 | 0.9793 |
| | | Juvenile | 0.06 ± 0.005 | 14 | | |
| | | Juvenilized-40 | 0.06 ± 0.004 | 20 | | |
| | | Juvenilized-78 | 0.06 ± 0.004 | 20 | | |
| | *A. thaliana* | Adult | 0.05 ± 0.031 | 6 | 2 | 0.3416 |
| | | Juvenile | 0.12 ± 0.029 | 7 | | |
| | | Juvenilized | 0.11 ± 0.029 | 7 | | |
| | *Zea mays* | Adult | 0.06 ± 0.002 a | 34 | 2 | <0.05 |
|  |  | Juvenile | 0.07 ± 0.002 ab | 38 |  |  |
|  |  | Juvenilized | 0.07 ± 0.002 b | 32 |  |  |
| LCP<br>($\mu\text{mol m}^{-2} \text{s}^{-1}$) | *P. tremula x alba* | Adult | 9.91 ± 1.23 a | 19 | 3 | <0.05 |
|  |  | Juvenile | 9.80 ± 1.43 a | 14 |  |  |
|  |  | Juvenilized-40 | 5.09 ± 1.26 b | 20 |  |  |
|  |  | Juvenilized-78 | 5.94 ± 1.20 b | 20 |  |  |
|  | *A. thaliana* | Adult | 5.10 ± 5.37 | 6 | 2 | 0.2395 |
|  |  | Juvenile | 16.21 ± 5.37 | 7 |  |  |
|  |  | Juvenilized | 16.90 ± 4.97 | 7 |  |  |
|  | *Zea mays* | Adult | 15.59 ± 1.12 | 34 | 2 | 0.1719 |
|  |  | Juvenile | 12.72 ± 1.06 | 38 |  |  |
|  |  | Juvenilized | 14.58 ± 1.16 | 32 |  |  |
| Fv/Fm | *P. tremula x alba* | Adult | 0.79 ± 0.010 a | 20 | 3 | <0.01 |
|  |  | Juvenile | 0.74 ± 0.012 b | 14 |  |  |
|  |  | Juvenilized-40 | 0.75 ± 0.010 b | 20 |  |  |
|  |  | Juvenilized-78 | 0.76 ± 0.010 b | 20 |  |  |
|  | *A. thaliana* | Adult | 0.77 ± 0.007 | 6 | 2 | 0.6823 |
|  |  | Juvenile | 0.77 ± 0.007 | 7 |  |  |
|  |  | Juvenilized | 0.76 ± 0.007 | 7 |  |  |
|  | *Zea mays* | Adult | 0.76 ± 0.004 | 31 | 2 | 0.4691 |
|  |  | Juvenile | 0.75 ± 0.004 | 38 |  |  |
|  |  | Juvenilized | 0.76 ± 0.004 | 31 |  |  |
| Fv'/Fm' | *P. tremula x alba* | Adult | 0.47 ± 0.014 | 19 | 3 | 0.4863 |
|  |  | Juvenile | 0.46 ± 0.016 | 14 |  |  |
|  |  | Juvenilized-40 | 0.44 ± 0.014 | 19 |  |  |
|  |  | Juvenilized-78 | 0.45 ± 0.013 | 19 |  |  |
|  | *A. thaliana* | Adult | 0.51 ± 0.010 | 6 | 2 | 0.2063 |
|  |  | Juvenile | 0.51 ± 0.009 | 7 |  |  |
|  |  | Juvenilized | 0.53 ± 0.009 | 7 |  |  |
|  | *Zea mays* | Adult | 0.47 ± 0.009 a | 31 | 2 | <0.01 |
|  |  | Juvenile | 0.44 ± 0.008 b | 38 |  |  |
|  |  | Juvenilized | 0.47 ± 0.009 a | 32 |  |  |
| $\Phi$ PSII | *P. tremula x alba* | Adult | 0.13 ± 0.008 a | 19 | 3 | <0.01 |
|  |  | Juvenile | 0.12 ± 0.009 ab | 14 |  |  |
|  |  | Juvenilized-40 | 0.09 ± 0.008 c | 19 |  |  |
|  |  | Juvenilized-78 | 0.11 ± 0.008 bc | 21 |  |  |
|  | *A. thaliana* | Adult | 0.11 ± 0.012 | 6 | 2 | 0.1825 |
|  |  | Juvenile | 0.08 ± 0.011 | 7 |  |  |
|  |  | Juvenilized | 0.09 ± 0.011 | 7 |  |  |
|  | *Zea mays* | Adult | 0.17 ± 0.007 | 31 | 2 | 0.3094 |
|  |  | Juvenile | 0.17 ± 0.006 | 38 |  |  |
|  |  | Juvenilized | 0.18 ± 0.007 | 32 |  |  |
| NPQ | *P. tremula x alba* | Adult | 3.42 ± 0.84 | 19 | 3 | 0.2044 |
|  |  | Juvenile | 5.30 ± 0.97 | 14 |  |  |
|  |  | Juvenilized-40 | 4.72 ± 0.84 | 19 |  |  |
|  |  | Juvenilized-78 | 5.84 ± 0.80 | 21 |  |  |
|  | *A. thaliana* | Adult | 5.76 ± 0.90 a | 6 | 2 | <0.01 |
|  |  | Juvenile | 0.43 ± 0.90 b | 7 |  |  |
|  |  | Juvenilized | 3.92 ± 0.84 a | 7 |  |  |
|  | *Zea mays* | Adult | 3.01 ± 0.54 | 31 | 2 | 0.6679 |
|  |  | Juvenile | 2.91 ± 0.49 | 38 |  |  |
|  |  | Juvenilized | 3.53 ± 0.53 | 32 |  |  |
| ETR<br>($\mu\text{mol m}^{-2} \text{s}^{-1}$) | *P. tremula x alba* | Adult | 87.45 ± 5.14 a | 19 | 3 | <0.01 |
|  |  | Juvenile | 81.91 ± 5.98 a | 14 |  |  |
|  |  | Juvenalized-40 | 60.08 ± 5.14 b | 19 |  |  |
|  |  | Juvenalized-78 | 69.60 ± 4.89 ab | 21 |  |  |
|  | *A. thaliana* | Adult | 48.15 ± 5.14 | 6 | 2 | 0.2081 |
|  |  | Juvenile | 35.17 ± 4.76 | 7 |  |  |
|  |  | Juvenalized | 40.45 ± 4.76 | 7 |  |  |
|  | *Zea mays* | Adult | 135.18 ± 5.42 | 31 | 2 | 0.3184 |
|  |  | Juvenile | 132.85 ± 4.89 | 38 |  |  |
|  |  | Juvenalized | 143.49 ± 5.33 | 32 |  |  |
| $R_{day}$<br>($\mu\text{mol m}^{-2} \text{s}^{-1}$) | *P. tremula x alba* | Adult | -0.78 ± 0.17 | 21 | 3 | 0.9685 |
|  |  | Juvenile | -0.77 ± 0.21 | 14 |  |  |
|  |  | Juvenalized-40 | -0.84 ± 0.18 | 19 |  |  |
|  |  | Juvenalized-78 | -0.89 ± 0.18 | 20 |  |  |
|  | *A. thaliana* | Adult | -1.20 ± 0.62 | 6 | 2 | 0.3251 |
|  |  | Juvenile | -1.41 ± 0.57 | 7 |  |  |
|  |  | Juvenalized | -0.23 ± 0.57 | 7 |  |  |
|  | *Zea mays* | Adult | -0.87 ± 0.16 | 30 | 2 | 0.184 |
|  |  | Juvenile | -0.43 ± 0.20 |  |  |  |
|  |  | Juvenalized | -0.84 ± 0.16 | 32 |  |  |
| $g_s \text{ High Light}$<br>($\text{mol m}^{-2} \text{s}^{-1}$) | *P. tremula x alba* | Adult | 0.195 ± 0.016 a | 19 | 3 | <0.0001 |
|  |  | Juvenile | 0.200 ± 0.018 a | 14 |  |  |
|  |  | Juvenalized-40 | 0.087 ± 0.087 b | 20 |  |  |
|  |  | Juvenalized-78 | 0.083 ± 0.015 b | 20 |  |  |
|  | *A. thaliana* | Adult | 0.103 ± 0.010 | 6 | 2 | 0.0869 |
|  |  | Juvenile | 0.060 ± 0.030 | 7 |  |  |
|  |  | Juvenalized | 0.098 ± 0.009 | 7 |  |  |
|  | *Zea mays* | Adult | 0.162 ± 0.009 | 34 | 2 | 0.3752 |
|  |  | Juvenile | 0.178 ± 0.007 | 38 |  |  |
|  |  | Juvenalized | 0.173 ± 0.009 | 32 |  |  |
| $g_s \text{ Low Light}$<br>($\text{mol m}^{-2} \text{s}^{-1}$) | *P. tremula x alba* | Adult | 0.010 ± 0.016 a | 19 | 3 | <0.05 |
|  |  | Juvenile | 0.156 ± 0.200 b | 14 |  |  |
|  |  | Juvenalized-40 | 0.068 ± 0.087 a | 20 |  |  |
|  |  | Juvenalized-78 | 0.073 ± 0.083 a | 20 |  |  |
|  | *A. thaliana* | Adult | 0.032 ± 0.010 | 6 | 2 | 0.1938 |
|  |  | Juvenile | 0.030 ± 0.010 | 7 |  |  |
|  |  | Juvenalized | 0.052 ± 0.009 | 7 |  |  |
|  | *Zea mays* | Adult | 0.042 ± 0.005 a | 34 | 2 | <0.0001 |
|  |  | Juvenile | 0.064 ± 0.004 b | 38 |  |  |
|  |  | Juvenalized | 0.042 ± 0.005 a | 32 |  |  |

**Table 3.** Photosynthetic nitrogen use efficiency (PNUE) represented by the slope of the linear relationship between A_max_ mass and leaf nitrogen. Variation in leaf nitrogen was too low to find any relationship in *A. thaliana*. *P*-values determined by least squares linear regression analysis across all leaves and by ANCOVA with developmental stage as the covariate.

| Species | Effect | Slope | y-intercept | $r^2$ | *p*-value |
| --- | --- | --- | --- | --- | --- |
| *P. tremula x alba* | All Stages | 0.25 | -0.30 | 0.55 | <0.0001 |
| | $N_{\text{mass}} \times \text{Developmental Stage}$ | | | | 0.8654 |
| *A. thaliana* | All Stages | *n.s.* | *n.s.* | *n.s.* | *n.s.* |
| | $N_{\text{mass}} \times \text{Developmental Stage}$ | | | | *n.s.* |
| *Zea mays* | All Stages | 0.42 | -0.11 | 0.50 | <0.0001 |
| | $N_{\text{mass}} \times \text{Developmental Stage}$ | | | | 0.1626 |

There were no significant differences in stomatal conductance (g_s_) or daytime respiration (R_d_) between adult and juvenile or juvenilized leaves in any of the test species (Table 2).

To determine if phase-dependent variation in A_max_ is attributable to variation in the biochemistry of photosynthesis, we examined traits modeled from AC_i_ curves (maximum Rubisco carboxylation rate, V_cmax_, and maximum electron transport rate for RuBP regeneration, J_max_), traits modeled from light response curves (quantum yield, Φ and light compensation point, LCP), and traits modeled from dark and light-adapted fluorescence (maximum quantum efficiency of PSII, Fv/Fm; maximum operating efficiency, Fv’/Fm’; quantum yield of photosystem II, ΦPSII; non-photochemical quenching, NPQ; and electron transport rate, ETR). With one exception, none of these traits were significantly different between adult vs. juvenile/juvenilized leaves. The sole exception was Fv/Fm in *P. tremula x alba*, which was 6.3% higher in adult leaves than juvenile leaves (Table 2).

The observation that phase-dependent variation in A_max_ is correlated with SLA and leaf N but not with most measures of photosynthetic or physiological efficiency suggests that phase-dependent aspects of leaf anatomy, as well as phase-dependent variation in leaf composition (e.g. protein content), are the primary determinants of variation in the rate of photosynthesis during shoot development.

### Low light photosynthetic traits

Under low light conditions (≤ 50 μmol m^-2^ s^-1^), adult and juvenile/juvenilized leaves of *P. tremula x alba* and *A. thaliana* showed no differences in area-based photosynthetic rates, whereas adult leaves of *Z. mays* had a slightly, but significantly lower A_max_ Area than juvenile or juvenilized leaves (Fig. **4**). This is in contrast to the relative rates of photosynthesis we observed at saturating light levels, where adult leaves of *P. tremula x alba* and *A. thaliana* had a significantly higher A_max_ Area than juvenile leaves, and the A_max_ Area in maize was not significantly different in these leaf types. The relative advantage of juvenile leaves under low light conditions was even more pronounced when photosynthesis was measured on a mass basis: in low light, juvenile and juvenilized leaves of all three species had a significantly higher A_max_ Mass than adult leaves. These results suggest that juvenile leaves are better adapted for photosynthesis under low light conditions than adult leaves.

**Figure 4.**
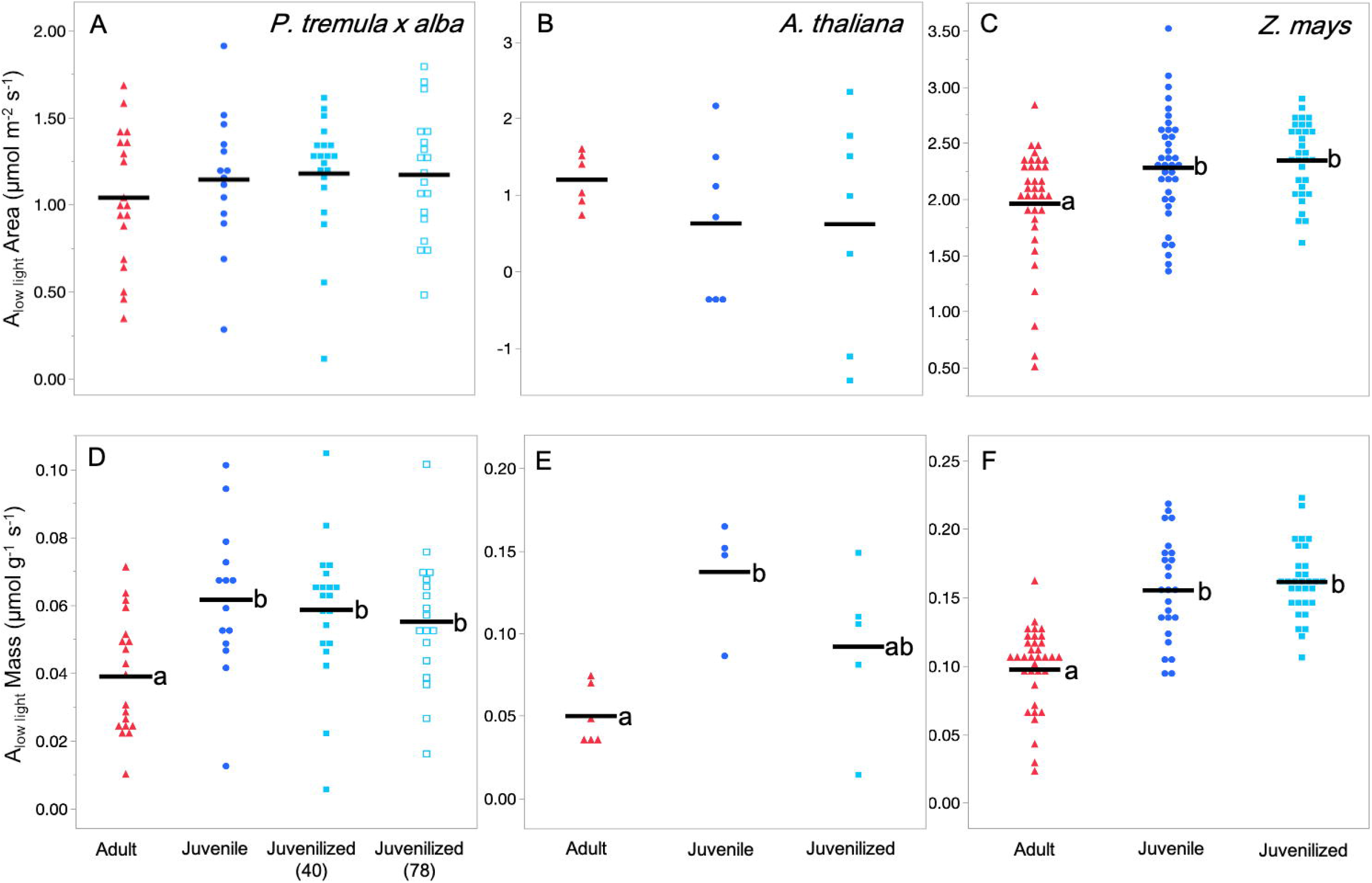
Low light photosynthetic rates for *P. tremula x alba* (A, D), *A. thaliana* (B, E) and *Z. mays* (C, F). Light levels were approximately 2-3x light compensation point at 25 μmol m^-2^ s^-1^ for *P. tremula x alba* and *A. thaliana* or 50 μmol m^-2^ s^-1^ for *Z. mays*. Traits depicted are area-based net photosynthetic rate at low light (A-C, A_low light_ Area) and mass-based net photosynthetic rate at low light (D-F, A_low light_ mass). Lettering and symbols are the same as Figure 1.

### Role of leaf position on phase-dependent traits

To determine whether there was an effect of leaf position—independent of phase identity— on various traits we looked across all measured leaf positions in wild-type and miR156 overexpressors of *P. tremula x alba* and *Z. mays*. Traits that varied with leaf number, but were not significantly different between wildtype and mutant plants were considered to be affected by leaf position independently of their phase identity. This is because wild-type plants had juvenile leaves at low nodes and adult leaves at high nodes, whereas miR156 overexpressors had juvenile leaves at all nodes. The only trait that showed a leaf position effect was A_low light_ Area in *Z. mays*, where values decreased with increasing leaf position regardless of developmental phase (Table 1).

### Photosynthetic traits in P. tremula x alba grown from seed

The analyses of *P. tremula x alba* described above were conducted with cuttings of the 717-1B4 clone propagated *in vitro*. We considered the first-formed leaves on these plants to be juvenile leaves because they differed morphologically from later-formed leaves, and because the leaves of transgenic plants over-expressing miR156 closely resembled these first-formed leaves. To determine how closely these plants resemble normal *P. tremula x alba*, we examined a variety of traits in successive leaves of plants grown from seeds. Consistent with the results obtained with plants propagated *in vitro*, SLA, A_max_ area, A_low_ area and Fv/Fm all showed significant differences between juvenile and adult leaves (Table 4). All other gas exchange and fluorescence traits did not display phase-specific differences, consistent with the results we obtained with 717-1B4 plants. These results demonstrate that vegetative phase change in *P. tremula x alba* plants regenerated *in vitro* is similar, if not identical, to vegetative phase change in seed-derived plants.

**Table 4.** Photosynthetic and leaf morphological traits for juvenile and adult leaves of *P. tremula x alba* grown from seed. *P*-values from one-way ANOVA with developmental stage or leaf position as the effect variable.

| Trait | Developmental Stage | Mean ± SE | N | Effect | df | *p*-value |
| --- | --- | --- | --- | --- | --- | --- |
| SLA<br>(cm<sup>2</sup> mg<sup>-1</sup>) | Adult | 0.20 ± 0.01 | 20 | Developmental Stage | 1 | <0.001 |
| | Juvenile | 0.25 ± 0.01 | 34 | Leaf Position | 1 | <0.001 |
| *A*<sub>max</sub> Area<br>(μmol m<sup>-2</sup> s<sup>-1</sup>) | Adult | 13.81 ± 1.39 | 20 | Developmental Stage | 1 | <0.01 |
| | Juvenile | 10.32 ± 0.91 | 39 | Leaf Position | 1 | <0.01 |
| *A*<sub>max</sub> Mass<br>(μmol g<sup>-2</sup> s<sup>-1</sup>) | Adult | 0.026 ± 0.005 | 20 | Developmental Stage | 1 | 0.4544 |
| | Juvenile | 0.020 ± 0.003 | 33 | Leaf Position | 1 | 0.0704 |
| *V*<sub>cmax</sub><br>(μmol m<sup>-2</sup> s<sup>-1</sup>) | Adult | 52.48 ± 5.89 | 20 | Developmental Stage | 1 | 0.1700 |
| | Juvenile | 35.71 ± 4.04 | 37 | Leaf Position | 1 | <0.001 |
| *J*<sub>max</sub><br>(μmol m<sup>-2</sup> s<sup>-1</sup>) | Adult | 84.34 ± 9.31 | 20 | Developmental Stage | 1 | 0.1959 |
| | Juvenile | 56.91 ± 6.39 | 37 | Leaf Position | 1 | <0.001 |
| *g*<sub>s</sub> High Light<br>(mol m<sup>-2</sup> s<sup>-1</sup>) | Adult | 0.206 ± 0.020 | 20 | Developmental Stage | 1 | 0.2695 |
| | Juvenile | 0.181 ± 0.014 | 39 | Leaf Position | 1 | 0.7735 |
| *A*<sub>low</sub> Area<br>(μmol m<sup>-2</sup> s<sup>-1</sup>) | Adult | 0.79 ± 0.079 | 20 | Developmental Stage | 1 | <0.05 |
| | Juvenile | 1.04 ± 0.057 | 39 | Leaf Position | 1 | 0.9176 |
| *A*<sub>low</sub> Mass<br>(μmol m<sup>-2</sup> s<sup>-1</sup>) | Adult | 0.002 ± 0.004 | 20 | Developmental Stage | 1 | 0.9623 |
| | Juvenile | 0.002 ± 0.003 | 33 | Leaf Position | 1 | 0.1190 |
| *g*<sub>s</sub> Low Light<br>(mol m<sup>-2</sup> s<sup>-1</sup>) | Adult | 0.102 ± 0.056 | 20 | Developmental Stage | 1 | 0.646 |
| | Juvenile | 0.139 ± 0.039 | 39 | Leaf Position | 1 | 0.9723 |
| Φ | Adult | 0.057 ± 0.006 | 20 | Developmental Stage | 1 | 0.2312 |
| | Juvenile | 0.053 ± 0.004 | 39 | Leaf Position | 1 | 0.0601 |
| LCP<br>(μmol m<sup>-2</sup> s<sup>-1</sup>) | Adult | 14.46 ± 2.00 | 20 | Developmental Stage | 1 | 0.585 |
| | Juvenile | 8.61 ± 1.31 | 39 | Leaf Position | 1 | 0.1317 |
| Fv/Fm | Adult | 0.78 ± 0.0156 | 20 | Developmental Stage | 1 | <0.01 |
| | Juvenile | 0.77 ± 0.010 | 39 | Leaf Position | 1 | <0.0001 |
| Φ PSII | Adult | 0.120 ± 0.020 | 20 | Developmental Stage | 1 | 0.0525 |
| | Juvenile | 0.092 ± 0.013 | 39 | Leaf Position | 1 | <0.01 |
| NPQ | Adult | 4.90 ± 1.74 | 20 | Developmental Stage | 1 | 0.4907 |
| | Juvenile | 5.75 ± 1.14 | 39 | Leaf Position | 1 | 0.6825 |
| ETR<br>(μmol m<sup>-2</sup> s<sup>-1</sup>) | Adult | 78.58 ± 12.91 | 20 | Developmental Stage | 1 | 0.0523 |
| | Juvenile | 60.48 ± 8.46 | 39 | Leaf Position | 1 | <0.01 |
| *R*<sub>day</sub><br>(μmol m<sup>-2</sup> s<sup>-1</sup>) | Adult | -1.09 ± 0.280 | 20 | Developmental Stage | 1 | 0.1667 |
| | Juvenile | -0.78 ± 0.184 | 39 | Leaf Position | 1 | <0.05 |

**Table 5.**
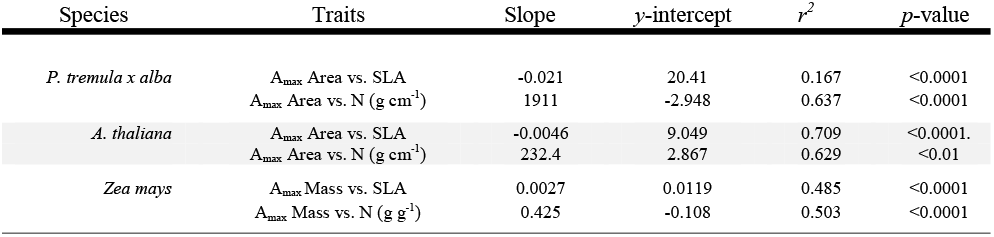
Linear fit between photosynthetic rates and leaf composition traits depicted in figure 3.

| Species | Traits | Slope | y-intercept | $r^2$ | $p$-value |
| --- | --- | --- | --- | --- | --- |
| *P. tremula x alba* | $A_{\max}$ Area vs. SLA | -0.021 | 20.41 | 0.167 | <0.0001 |
| | $A_{\max}$ Area vs. N ($\text{g cm}^{-1}$) | 1911 | -2.948 | 0.637 | <0.0001 |
| *A. thaliana* | $A_{\max}$ Area vs. SLA | -0.0046 | 9.049 | 0.709 | <0.0001 |
| | $A_{\max}$ Area vs. N ($\text{g cm}^{-1}$) | 232.4 | 2.867 | 0.629 | <0.01 |
| *Zea mays* | $A_{\max}$ Mass vs. SLA | 0.0027 | 0.0119 | 0.485 | <0.0001 |
| | $A_{\max}$ Mass vs. N ($\text{g g}^{-1}$) | 0.425 | -0.108 | 0.503 | <0.0001 |

**Table 6.**
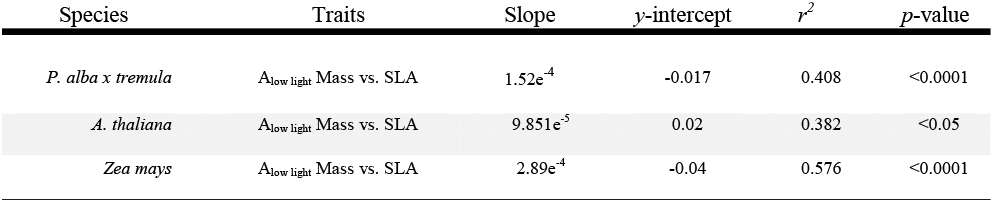
Linear fit between mass-based low light photosynthetic rates and SLA.

| Species | Traits | Slope | y-intercept | $r^2$ | $p$-value |
| --- | --- | --- | --- | --- | --- |
| *P. alba x tremula* | $A_{\text{low light}}$ Mass vs. SLA | $1.52\text{e}^{-4}$ | -0.017 | 0.408 | <0.0001 |
| *A. thaliana* | $A_{\text{low light}}$ Mass vs. SLA | $9.851\text{e}^{-5}$ | 0.02 | 0.382 | <0.05 |
| *Zea mays* | $A_{\text{low light}}$ Mass vs. SLA | $2.89\text{e}^{-4}$ | -0.04 | 0.576 | <0.0001 |

## Discussion

Numerous studies have shown that leaves produced at different times in plant development often have different rates of photosynthesis (Bond, 2000). Here, we investigated whether this phenomenon can be attributed to the transition between juvenile and adult phases of vegetative development, a process called vegetative phase change. Previous studies have described differences in photosynthetic efficiency between juvenile and adult leaves of strongly heteroblastic species of *Eucalyptus* (Cameron, 1970; Velikova *et al*., 2008) and *Acacia* (Brodribb & Hill, 1993; Hansen, 1996; Yu & Li, 2007). However, it is difficult to know if these studies are generally relevant because of the large anatomical differences between juvenile and adult leaves in these species, and because these studies did not control for the effect of leaf position. We characterized how vegetative phase change impacts photosynthesis independent of other confounding factors by manipulating the expression of miR156, the master regulator of this process. The miR156 overexpressors used in this study delay vegetative phase change, causing the plants to produce leaves with juvenile identity at positions that are normally adult. This made it possible to distinguish miR156-regulated photosynthetic traits from photosynthetic traits that vary as function of leaf position or plant age.

In all three of the species we examined (*P. tremula x alba, A. thaliana*, and *Z. mays*) the rate of light-saturated photosynthesis was phase-dependent, although this relationship differed between species depending on whether area- or mass-based measures were used. Previous studies have revealed significant differences in the expression of photosynthetic genes in juvenile and adult leaves of *Z. mays* (Strable *et al*., 2008; Beydler, 2014) and *Malus domestica* Borkh.(Gao *et al*., 2014), suggesting that phase-dependent variation in the rate of photosynthesis might be attributable to differences in the biochemistry of photosynthesis in different leaves. However, multiple measures of photosynthetic capacity and light use efficiency provided no evidence of this. Instead, we found that the difference in the rate of photosynthesis in juvenile and adult leaves was most highly correlated with differences in the SLA and N content of these leaves. This observation suggests that phase-dependent differences in photosynthetic rates are attributable to differences in leaf anatomy and leaf composition, rather than differences in the biochemistry of photosynthesis.

Leaf morphology and composition have robust relationships with photosynthesis across species and environments (Niinemets & Tenhunen, 1997; Reich *et al*., 1998, 1999, 2003; Meziane & Shipley, 2001). Leaf thickness and density—the structural changes that determine SLA— modulate intra-leaf light dynamics, CO_2_ diffusion and the distribution of leaf N (Parkhurst, 1994; Epron *et al*., 1995; Terashima & Hikosaka, 1995; Reich *et al*., 1998; Terashima *et al*., 2006; Evans *et al*., 2009). Specifically, variation in SLA changes the way light moves within the leaf as path length and scattering is altered. This leads to leaves with low SLA absorbing more light per area as pathlength increases, ultimately leading to higher A_max_ area (Terashima & Hikosaka, 1995). However, leaves with low SLA face the challenge of increased CO_2_ diffusion resistance as CO_2_ must travel farther from stomata and through denser tissue to reach carboxylating enzymes (Parkhurst, 1994; Terashima *et al*., 2006). SLA further impacts photosynthesis through the distribution of leaf N as leaves with low SLA are associated with more cytoplasmic volume per leaf area and therefore more N. The relationship between leaf N and photosynthesis results from the well-established relationship between N, Rubisco and other photosynthetically important proteins (Field & Mooney, 1986; Evans, 1989; Ellsworth & Reich, 1993; Makino *et al*., 1994; Bond *et al*., 1999; Chmura & Tjoelker, 2008).

It is currently unclear why phase-dependence in A_max_ and leaf N are observed in area-based measures for *P. tremula x alba* and *A. thaliana* but mass-based measures for *Z. mays* (although the fact that only one form of measurement correlates with SLA and leaf N is expected) (Westoby *et al*., 2013). These three species all have relatively high SLA, and no differences in PNUE between juvenile and adult leaves, which would suggest differences in the A_max_-N slope due to SLA (Reich *et al*., 1998) do not contribute to this phenomenon. Other potential explanations include differences in photosynthetic pathway (C_3_ vs. C_4_), developmental form (dicot vs. monocot) or variation in the morphological contributors to SLA (leaf thickness vs. cell density). Because the relationships between SLA and photosynthetic rate are conserved across data sets that include both C_3_ and C_4_ species as well as both monocots and dicots these traits are unlikely to explain the differences between species in this study (Reich *et al*., 1999, 2003; Meziane & Shipley, 2001). While density and thickness each contribute to variation in SLA, the degree to which they alter the photosynthetically important properties of a leaf vary. Because of this, Niinemets (1999) found that changes in leaf thickness are more closely correlated with area-based photosynthetic rates while changes in density with mass-based rates. As to be expected, changes in both leaf thickness and density have been associated with changes in SLA across all three study species (Bongard-Pierce *et al*., 1996; Wang *et al*., 2011; Chuck *et al*., 2011; Coneva & Chitwood, 2018) and can be observed in cross sections of adult, juvenile and juvenilized leaves in this study (Fig. **5**). Further studies are needed to determine the extent to which density and thickness contribute to phase-dependent changes in SLA and the mass or area-based correlations observed in this study.

**Figure 5.**
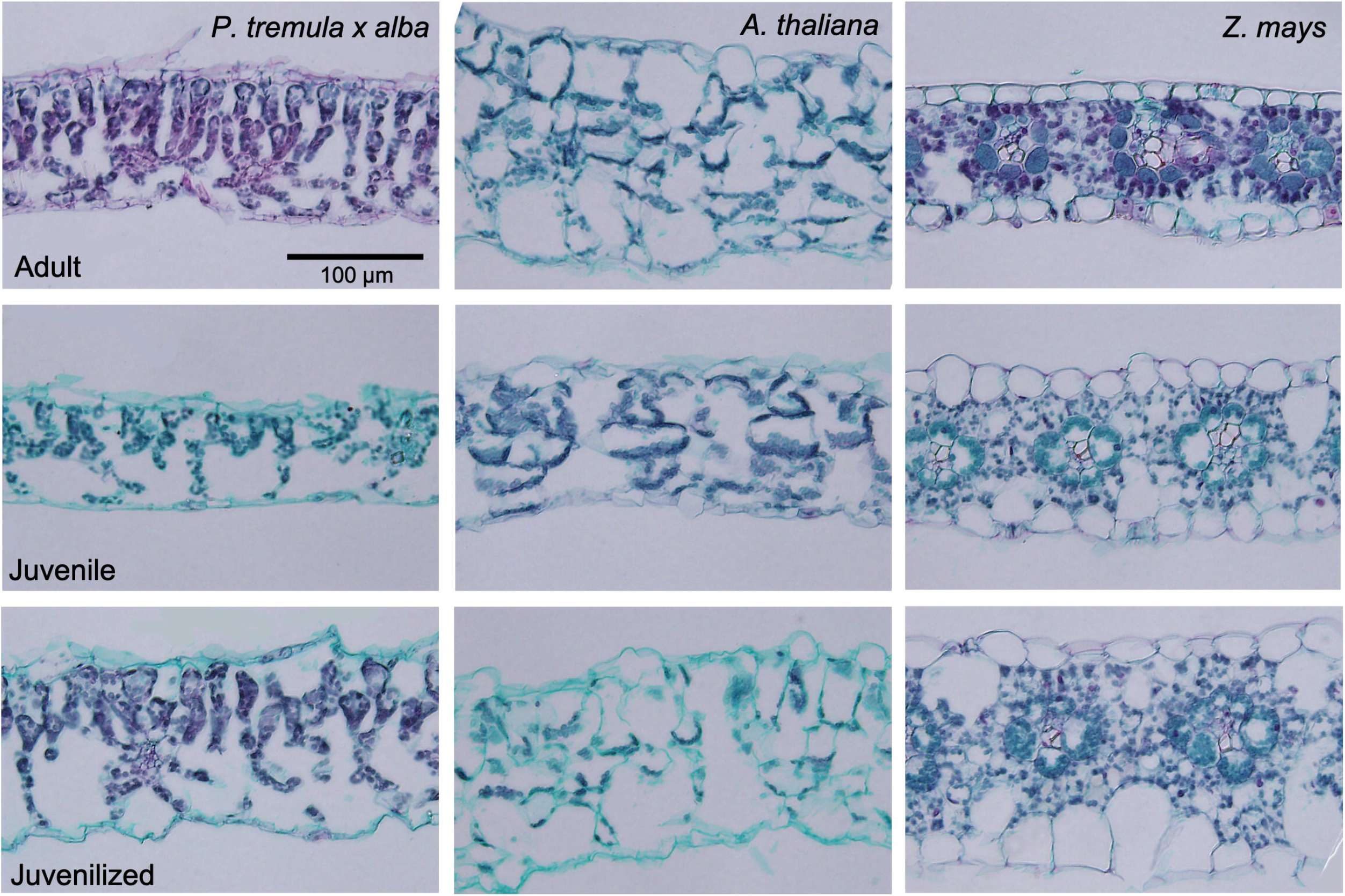
Leaf cross sections of *P. tremula x alba, A. thaliana*, and *Z. mays* adult, juvenile and juvenilized leaves stained with safranin-O and fast green.

Juvenile leaf morphology and photosynthetic properties may contribute to better survival in low light environments, such as those frequently experienced by juvenile tissues at the bottom of a canopy. High SLA, found in juvenile leaves of all three species, is strongly correlated with higher photosynthetic rates under light limited conditions and shade-tolerance (Givnish, 1988; Niinemets & Tenhunen, 1997; Walters & Reich, 1999; Reich *et al*., 2003). In support of this hypothesis, the juvenile leaves in each species had higher mass-based photosynthetic rates at low light levels (A_low light_) than adult leaves. Even in area-based measures of *P. tremula x alba* and *A. thaliana*, where adult leaves have higher A_max_, this photosynthetic advantage is lost under light-limited conditions. Further, variation in photosynthesis and SLA have been associated with tolerance to additional environmental factors, including drought and herbivory, and with changes in growth strategy such as leaf life-span and growth rate (Poorter, 1999; Wright & Cannon, 2001; Reich *et al*., 2003; Poorter *et al*., 2009; Niinemets, 2010; Dayrell *et al*., 2018). Because these traits are phase-dependent, it is likely vegetative phase change contributes to variation in biotic and abiotic stress tolerance during a plant’s lifetime.

The broad documentation of decreasing SLA and photosynthetic variation during plant growth suggests the phase-dependence of these traits goes beyond the species examined here. Further, this study provides evidence that miR156 and the regulators of phase change are an endogenous mechanism contributing to the developmental variation in these traits independent of plant size and age. Because of its role in leaf morphology and photosynthetic properties, the timing of VPC could have important implications for selection and adaptation as climates change globally. While more studies are needed regarding this topic, vegetative phase change has the potential to contribute significantly to species adaptation and acclimation during plant vegetative growth.

## Acknowledgements

We thank Samara Gray and Joshua Darfler for their assistance in caring for the plants used in this study and Che-Ling Ho for assistance with leaf cross sections. This research was funded by the NSF Graduate Research Fellowship (Division of Graduate Education; DGE-1321851), U. of Pennsylvania SAS Dissertation Research Fellowship and the Peachey Research Fund awarded to E.H.L.

## Author Contributions

E.H.L., C.J.S., B.R.H., and R.S.P. planned and designed the research. E.H.L. performed the experiments. E.H.L performed statistical analyses and wrote the manuscript. E.H.L., C.J.S., B.R.H., and R.S.P revised and provided comments on the manuscript.

## References

Ahsan MU, Hayward A, Alam M, Bandaralage JH, Topp B, Beveridge CA, Mitter N. 2019. Scion control of miRNA abundance and tree maturity in grafted avocado. BMC Plant Biology 19: 382.

Axtell MJ, Bowman JL. 2008. Evolution of plant microRNAs and their targets. Trends in Plant Science 13: 343–349.

Bassiri A, Irish EE, Scott PR. 1992. Heterochronic effects of Teopod 2 on the growth and photosensitivity of the maize shoot. The Plant Cell 4: 497–504.

Bauer H, Bauer U. 1980. Photosynthesis in leaves of the juvenile and adult phase of ivy (Hedera helix). Physiologia plantarum 49: 366–372.

Beydler BD. 2014. Dynamics of gene expression during vegetative phase change in dynamics of gene expression during vegetative phase change in maize. Ph.D. Thesis.

Bhogale S, Mahajan AS, Natarajan B, Rajabhoj M, Thulasiram H V, Banerjee AK. 2014. MicroRNA156: A potential graft-transmissible microrna that modulates plant architecture and tuberization in Solanum tuberosum ssp. andigena. Plant Physiology 164: 1011–1027.

Bond BJ. 2000. Age-related changes in photosynthesis of woody plants. Trends in Plant Science 5: 349–353.

Bond BJ, Farnsworth BT, Coulombe RA, Winner WE. 1999. Foliage physiology and biochemistry in response to light gradients in conifers with varying shade tolerance. Oecologia 120:183–192.

Bongard-Pierce DK, Evans MMS, Poethig RS. 1996. Heteroblastic features of leaf anatomy in maize and their genetic regulation. International Journal of Plant Sciences 157: 331.

Brodribb T, Hill RS. 1993. A physiological comparison of leaves and phyllodes in Acacia melanoxylon. Australian Journal of Botany 41: 293–305.

Cameron RJ. 1970. Light intensity and the growth of Eucalyptus seedlings. I. Ontogenetic variation in E. Fastigata. Australian Journal of Botany 18: 29–43.

Canham CD. 1988. An index for understory light levels in and around canopy gaps. Ecology 69: 1634–1638.

Canham CD, Finzi AC, Pacala SW, Burbank DH. 1994. Causes and consequences of resource heterogeneity in forests: Interspecific variation in light transmission by canopy trees. Canadian Journal of Forest Research 24: 337–349.

Cavender-Bares J, Bazzaz FA. 2000. Changes in drought response strategies with ontogeny in Quercus rubra: implications for scaling from seedlings to mature trees. Oecologia 124: 8–18.

Charles LS, Dwyer JM, Smith TJ, Connors S, Marschner P, Mayfield MM. 2018. Seedling growth responses to species-, neighborhood-, and landscape-scale effects during tropical forest restoration. Ecosphere 9: e02386.

Chmura DJ, Tjoelker MG. 2008. Leaf traits in relation to crown development, light interception and growth of elite families of loblolly and slash pine. Tree Physiology 28: 729–742.

Chuck G, Cigan M, Saeteurn K, Hake S. 2007. The heterochronic maize mutant Corngrass1 results from overexpression of a tandem microRNA. Nature genetics 39: 544–549.

Chuck GS, Tobias C, Sun L, Kraemer F, Li C, Dibble D, Arora R, Bragg JN, Vogel JP, Singh S, et al. 2011. Overexpression of the maize Corngrass1 microRNA prevents flowering, improves digestibility, and increases starch content of switchgrass. Proceedings of the National Academy of Sciences 109: 995–995.

Coneva V, Chitwood DH. 2018. Genetic and developmental basis for increased leaf thickness in the arabidopsis cvi ecotype. Frontiers in Plant Science 9.

Dayrell RLC, Arruda AJ, Pierce S, Negreiros D, Meyer PB, Lambers H, Silveira FAO. 2018. Ontogenetic shifts in plant ecological strategies. Functional Ecology 32: 2730–2741.

Duursma RA. 2015. Plantecophys - An R package for analysing and modelling leaf gas exchange data. PLoS ONE 10: e0143346.

Ellsworth DS, Reich PB. 1993. Canopy structure and vertical patterns of photosynthesis and related leaf traits in a deciduous forest. Oecologia 96: 169–178.

Epron D, Godard D, Cornic G, Genty B. 1995. Limitation of net CO2 assimilation rate by internal resistances to CO2 transfer in the leaves of two tree species (Fagus sylvatica L. and Castanea sativa Mill.). Plant, Cell & Environment 18: 43–51.

Evans JR. 1989. Photosynthesis and nitrogen relationships in leaves of C□ plants. Oecologia 78: 9–19.

Evans GC, Coombe DE. 1959. Hemisperical and woodland canopy photography and the light climate. The Journal of Ecology 47: 103.

Evans JR, Kaldenhoff R, Genty B, Terashima I. 2009. Resistances along the CO2 diffusion pathway inside leaves. Journal of Experimental Botany 60: 2235–2248.

Feng S, Xu Y, Guo C, Zheng J, Zhou B, Zhang Y, Ding Y, Zhang L, Zhu Z, Wang H, et al. 2016. Modulation of miR156 to identify traits associated with vegetative phase change in tobacco (Nicotiana tabacum). Journal of Experimental Botany 67: 1493–1504.

Field C, Mooney HA. 1986. The photosynthesis-nitrogen relationship in wild plants. In: Givnish TJ, ed. On the Economy of Plant Form and Function. Cambridge University Press, 25–55.

Flexas J, Ortuño MF, Ribas-Carbo M, Diaz-Espejo A, Flórez-Sarasa ID, Medrano H. 2007. Mesophyll conductance to CO2 in Arabidopsis thaliana. New Phytologist 175: 501–511.

Fouracre JP, Poethig RS. 2019. Role for the shoot apical meristem in the specification of juvenile leaf identity in Arabidopsis. Proceedings of the National Academy of Sciences of the United States of America 116: 10168–10177.

Gao Y, Yang FQ, Cao X, Li CM, Wang Y, Zhao YB, Zeng GJ, Chen DM, Han ZH, Zhang XZ. 2014. Differences in gene expression and regulation during ontogenetic phase change in apple seedlings. Plant Molecular Biology Reporter 32: 357–371.

Givnish TJ. 1988. Adaptation to sun and shade: a whole-plant perspective. Australian Journal of Plant Physiology 15: 63–92.

Grossnickle SC. 2012. Why seedlings survive: influence of plant attributes. New Forests 43: 711–738.

Hahn PG, Orrock JL. 2016. Neighbor palatability generates associational effects by altering herbivore foraging behavior. Ecology 97: 2103–2111.

Hansen DH. 1996. Establishment and persistence characteristics in juvenile leaves and phyllodes of Acacia koa (leguminosae) in Hawaii. International Journal of Plant Sciences 157: 123–128.

He J, Xu M, Willmann MR, McCormick K, Hu T, Yang L, Starker CG, Voytas DF, Meyers BC, Poethig RS. 2018. Threshold-dependent repression of SPL gene expression by miR156/miR157 controls vegetative phase change in Arabidopsis thaliana. PLOS Genetics 14: e1007337.

Huang L-C, Weng J-H, Wang C-H, Kuo C-I, Shieh Y-J. 2003. Photosynthetic potentials of in vitro-grown juvenile, adult, and rejuvenated Sequoia sempervirens (D. Don) Endl. shoots. Botanical Bulletin of Academia Sinica 44: 31–35.

Hutchison KW, Sherman CD, Weber J, Smith SS, Singer PB, Greenwood MS. 1990. Maturation in Larch: II. Effects of age on photosynthesis and gene expression in developing foliage. Plant physiology 94: 1308–15.

Irish EE, Karlen S. 1998. Restoration of juvenility in maize shoots by meristem culture. International Journal of Plant Sciences 159: 695–701.

Ishida A, Yazaki K, Hoe AL. 2005. Ontogenetic transition of leaf physiology and anatomy from seedlings to mature trees of a rain forest pioneer tree, Macaranga gigantea. Tree Physiology 25:513–522.

Jaya E, Kubien DS, Jameson PE, Clemens J. 2010. Vegetative phase change and photosynthesis in Eucalyptus occidentalis: architectural simplification prolongs juvenile traits. Tree Physiology 30: 393–403.

Kabrick JM, Knapp BO, Dey DC, Larsen DR. 2015. Effect of initial seedling size, understory competition, and overstory density on the survival and growth of Pinus echinata seedlings underplanted in hardwood forests for restoration. New Forests 46: 897–918.

Kerr KL, Meinzer FC, McCulloh KA, Woodruff DR, Marias DE. 2015. Expression of functional traits during seedling establishment in two populations of Pinus ponderosa from contrasting climates. Tree Physiology 35: 535–548.

Kitajima K, Cordero RA, Wright SJ. 2013. Leaf life span spectrum of tropical woody seedlings: effects of light and ontogeny and consequences for survival. Annals of Botany 112: 685–699.

Kubien D, Jaya E, Clemens J. 2007. Differences in the structure and gas exchange physiology of juvenile and adult leaves in metrosideros excelsa. International Journal of Plant Sciences 168: 563–570.

Kuusk V, Niinemets Ü, Valladares F. 2018a. A major trade-off between structural and photosynthetic investments operative across plant and needle ages in three Mediterranean pines. Tree Physiology 38: 543–557.

Kuusk V, Niinemets Ü, Valladares F. 2018b. Structural controls on photosynthetic capacity through juvenile-to-adult transition and needle ageing in Mediterranean pines. Functional Ecology 32: 1479–1491.

Lamb EG, Cahill JF. 2006. Consequences of differing competitive abilities between juvenile and adult plants. Oikos 112: 502–512.

Lasky JR, Bachelot B, Muscarella R, Schwartz N, Forero-Montaña J, Nytch CJ, Swenson NG, Thompson J, Zimmerman JK, Uriarte M. 2015. Ontogenetic shifts in trait-mediated mechanisms of plant community assembly. Ecology 96: 2157–2169.

Lawrence EH, Leichty AR, Ma C, Strauss SH, Poethig RS. 2020. Vegetative phase change in Populus tremula x alba. bioRxiv: 2020.06.21.163469.

Lawrence EH, Stinziano JR, Hanson DT. 2019. Using the rapid A-C i response (RACiR) in the Li-Cor 6400 to measure developmental gradients of photosynthetic capacity in poplar. Plant Cell and Environment 42: 740–750.

Leichty AR, Poethig RS. 2019. Development and evolution of age-dependent defenses in ant-acacias. Proceedings of the National Academy of Sciences 116: 15596–15601.

Li H, Zhao X, Dai H, Wu W, Mao W, Zhang Z. 2012. Tissue culture responsive microRNAs in strawberry. Plant Molecular Biology Reporter 30: 1047–1054.

Lobo F de A, de Barros MP, Dalmagro HJ, Dalmolin ÂC, Pereira WE, de Souza ÉC, Vourlitis GL, Rodríguez Ortíz CE. 2013. Fitting net photosynthetic light-response curves with Microsoft Excel - a critical look at the models. Photosynthetica 51: 445–456.

Lusk CH, Del Pozo A. 2002. Survival and growth of seedlings of 12 Chilean rainforest trees in two light environments: Gas exchange and biomass distribution correlates. Austral Ecology 27: 173–182.

Makino A, Nakano H, Mae T. 1994. Responses of ribulose-1,5-bisphosphate carboxylase, cytochrome f, and sucrose synthesis enzymes in rice leaves to leaf nitrogen and their relationships to photosynthesis. Plant Physiology 105: 173–179.

Marin-Gonzalez E, Suarez-Lopez P. 2012. “And yet it moves”: Cell-to-cell and long-distance signaling by plant microRNAs. Plant Science 196: 18–30.

Meilan R, Ma C. 2006. Poplar (Populus spp.). In: Wang K, ed. Methods in Molecular Biology: Agrobacterium Protocols. Totowa, NJ: Humana Press Inc., 143–151.

Meziane D, Shipley B. 2001. Direct and indirect relationships between specific leaf area, leaf nitrogen and leaf gas exchange. Effects of irradiance and nutrient supply. Annals of Botany 88: 915–927.

Modrzynski J, Chmura DJ, Tjoelker MG. 2015. Seedling growth and biomass allocation in relation to leaf habit and shade tolerance among 10 temperate tree species. Tree Physiology 35: 879–893.

Moll JD, Brown JS. 2008. Competition and coexistence with multiple life□history stages. The American Naturalist 171: 839–843.

Niinemets Ü. 1999. Components of leaf dry mass per area - thickness and density - alter leaf photosynthetic capacity in reverse directions in woody plants. New Phytologist 144: 35–47.

Niinemets Ü. 2010. Responses of forest trees to single and multiple environmental stresses from seedlings to mature plants: Past stress history, stress interactions, tolerance and acclimation. Forest Ecology and Management 260: 1623–1639.

Niinemets Ü, Tenhunen JD. 1997. A model separating leaf structural and physiological effects on carbon gain along light gradients for the shade-tolerant species Acer saccharum. Plant, Cell and Environment 20: 845–866.

Parish JAD, Bazzaz FA. 1985. Ontogenetic niche shifts in old-field annuals. Ecology 66: 1296–1302.

Parkhurst DF. 1994. Diffusion of CO2 and other gases inside leaves. New Phytologist 126: 449–479.

Piao T, Comita LS, Jin G, Kim JH. 2013. Density dependence across multiple life stages in a temperate old-growth forest of northeast China. Oecologia 172: 207–217.

Poethig RS. 1988. Heterochronic mutations affecting shoot development in maize. Genetics 119: 959–73.

Poethig RS. 1990. Phase change and the regulation of shoot morphogenesis in plants. Science.

Poorter L. 1999. Growth responses of 15 rain-forest tree species to a light gradient: The relative importance of morphological and physiological traits. Functional Ecology 13: 396–410.

Poorter L, Bongers F. 2006. Leaf traits are good predictors of plant performance across 53 rain forest species. Ecology 87: 1733–1743.

Poorter H, Niinemets Ü, Poorter L, Wright IJ, Villar R. 2009. Causes and consequences of variation in leaf mass per area (LMA): a meta analysis. New Phytologist 182: 565–588.

Porra RJ, Thompson WA, Kriedemann PE. 1989. Determination of accurate extinction coefficients and simultaneous equations for assaying chlorophylls a and b extracted with four different solvents: verification of the concentration of chlorophyll standards by atomic absorption spectroscopy. Biochimica et Biophysica Acta 975: 384–394.

Reich PB, Ellsworth DS, Walters MB. 1998. Leaf structure (specific leaf area) modulates photosynthesis–nitrogen relations: evidence from within and across species and functional groups. Functional Ecology 12: 948–958.

Reich PB, Ellsworth DS, Walters MB, Vose JM, Gresham C, Volin JC, Bowman WD. 1999. Generality of leaf trait relationships: A test across six biomes. Ecology 80: 1955–1969.

Reich PB, Wright IJ, Cavender-Bares J, Craine JM, Oleksyn J, Westoby M, Walters MB. 2003. The evolution of plant functional variation: Traits, spectra, and strategies. International Journal of Plant Sciences 164: S143–S164.

Shuttleworth WJ, Gash JHC, Lloyd CR, Moore CJ, Roberts J, Marques Filho A de O, Fisch G, Silva Filho V de P, Nazare Goes Ribeiro M de, Molion LCB, et al. 1985. Daily variations of temperature and humidity within and above amazonian forest. Weather 40: 102–108.

Silva PO, Batista DS, Henrique J, Cavalcanti F, Koehler AD, Vieira LM, Fernandes AM, Hernan Barrera-Rojas C, Ribeiro DM, Nogueira FTS, et al. 2019. Leaf heteroblasty in Passiflora edulis as revealed by metabolic profiling and expression analyses of the microRNAs miR156 and miR172. Annals of Botany 123: 1191–1203.

Spasojevic MJ, Yablon EA, Oberle B, Myers JA. 2014. Ontogenetic trait variation influences tree community assembly across environmental gradients. Ecosphere 5: art129.

Steppe K, Niinemets Ü, Teskey RO. 2011. Tree size- and age-related changes in leaf physiology and their influence on carbon gain. In: Meinzer FC, Lachenbruch B, Dawson TE, eds. Size- and age-related changes in tree structure and function. Springer, 235–253.

Still C, Powell R, Aubrecht D, Kim Y, Helliker B, Roberts D, Richardson AD, Goulden M. 2019. Thermal imaging in plant and ecosystem ecology: applications and challenges. Ecosphere 10: e02768.

Strable J, Borsuk L, Nettleton D, Schnable PS, Irish EE. 2008. Microarray analysis of vegetative phase change in maize. Plant Journal 56: 1045–1057.

Sun J, Yao F, Wu J, Zhang P, Xu W. 2018. Effect of nitrogen levels on photosynthetic parameters, morphological and chemical characters of saplings and trees in a temperate forest. Journal of Forestry Research 29: 1481–1488.

Telfer A, Bollman KM, Poethig RS. 1997. Phase change and the regulation of trichome distribution in Arabidopsis thaliana. Development 124: 645–654.

Terashima I, Hanba YT, Tazoe Y, Vyas P, Yano S. 2006. Irradiance and phenotype: Comparative eco-development of sun and shade leaves in relation to photosynthetic CO2 diffusion. In: Journal of Experimental Botany. 343–354.

Terashima I, Hikosaka K. 1995. Comparative ecophysiology of leaf and canopy photosynthesis. Plant, Cell & Environment 18: 1111–1128.

Tomeo N. 2019. Tomeopaste/AQ_curves: AQ_curve fitting release 1.

Velikova V, Loreto F, Brilli F, Stefanov D, Yordanov I. 2008. Characterization of juvenile and adult leaves of Eucalyptus globulus showing distinct heteroblastic development: photosynthesis and volatile isoprenoids. Plant Biology 10: 55–64.

Waggoner PE, Reifsnyder WE. 1968. Simulation of the temperature, humidity and evaporation profiles in a leaf canopy. Journal of Applied Meteorology 7: 400–409.

Walters MB, Reich PB. 1999. Low-light carbon balance and shade tolerance in the seedlings of woody plants: Do winter deciduous and broad-leaved evergreen species differ? New Phytologist 143:143–154.

Wang JW, Park MY, Wang LJ, Koo Y, Chen XY, Weigel D, Poethig RS. 2011. MiRNA control of vegetative phase change in trees. PLoS Genetics 7: e1002012.

Westoby M, Reich PB, Wright IJ. 2013. Understanding ecological variation across species: Area-based vs mass-based expression of leaf traits. New Phytologist 199: 322–323.

Willmann MR, Poethig RS. 2007. Conservation and evolution of miRNA regulatory programs in plant development. Current Opinion in Plant Biology 10: 503–511.

Wright IJ, Cannon K. 2001. Relationships between leaf lifespan and structural defences in a low-nutrient, sclerophyll flora. Functional Ecology 15: 351–359.

Wu G, Park MY, Conway SR, Wang JW, Weigel D, Poethig RS. 2009. The sequential action of miR156 and miR172 regulates developmental timing in Arabidopsis. Cell 138: 750–759.

Wu G, Poethig RS. 2006. Temporal regulation of shoot development in Arabidopsis thaliana by miR156 and its target SPL3. Development 133: 3539–3547.

Xu M, Hu T, Zhao J, Park M-Y, Earley KW, Wu G, Yang L, Poethig RS. 2016. Developmental functions of miR156-regulated SQUAMOSA PROMOTER BINDING PROTEIN-LIKE (SPL) genes in arabidopsis thaliana. PLOS Genetics 12: e1006263.

Yin X, Sun Z, Struik PC, Van Der Putten PEL, Van Ieperen W, Harbinson J. 2011. Using a biochemical C4 photosynthesis model and combined gas exchange and chlorophyll fluorescence measurements to estimate bundle-sheath conductance of maize leaves differing in age and nitrogen content. Plant, Cell and Environment 34: 2183–2199.

Yu H, Li JT. 2007. Physiological comparisons of true leaves and phyllodes in Acacia mangium seedlings. Photosynthetica 45: 312–316.

Zhang SD, Ling LZ, Zhang QF, Xu J Di, Cheng L. 2015. Evolutionary comparison of two combinatorial regulators of SBP-Box genes, miR156 and miR529, in plants. PLoS ONE 10: e0124621.

Zhou H, Akçay E, Helliker BR. 2019. Estimating C4 photosynthesis parameters by fitting intensive A/Ci curves. Photosynthesis Research 141: 181–194.

